# Nickel induced transcriptional changes persist post exposure through epigenetic reprograming

**DOI:** 10.1101/806588

**Authors:** Cynthia C Jose, Zhenjia Wang, Vinay Singh Tanwar, Xiaoru Zhang, Chongzhi Zang, Suresh Cuddapah

**Affiliations:** Department of Environmental Medicine, New York University School of Medicine, New York, NY 10010, USA; Center for Public Health Genomics, Department of Public Health Sciences, University of Virginia School of Medicine, Charlottesville, VA 22908, USA

**Keywords:** Nickel exposure, environmental toxicant, post-exposure epigenome changes, persistent gene differential expression, epigenome, H3K4me3, H3K27me3

## Abstract

Nickel is an occupational and environmental toxicant associated with a number of diseases in humans including pulmonary fibrosis, bronchitis and lung and nasal cancers. Our earlier studies showed that the nickel-exposure-induced genome-wide transcriptional changes, which persist even after the termination of exposure may underlie nickel pathogenesis. However, the mechanisms that drive nickel-induced persistent changes to the transcriptome remain elusive. To elucidate the mechanisms that underlie nickel induced long-term transcriptional changes, in this study, we examined the transcriptome and the epigenome of human lung epithelial cells during nickel exposure and after the termination of exposure. We identified two categories of persistently differentially expressed genes based on the timing of expression changes: i) the genes that were differentially expressed during nickel exposure; and ii) the genes that were differentially expressed only after the termination of nickel exposure. Interestingly, the majority of nickel-induced transcriptional changes occurred only after the termination of exposure. We found robust genome-wide alterations to the activating histone modification, H3K4me3, after the termination of nickel exposure, which coincided with the post-exposure gene expression changes. In addition, we found significant post-exposure alterations to the repressive histone modification, H3K27me3. By uncovering a new category of transcriptional and epigenetic changes, which occur only after the termination of exposure, this study sheds new light on the post-exposure effects of nickel and provides a novel understanding of the long-term deleterious consequences of nickel exposure on human health.

## Introduction

Nickel is an environmental and occupational toxicant and exposure to nickel is associated with multiple diseases in humans including lung and nasal cancers [1–3]. While previous studies have examined the deleterious effects of nickel on a number of cellular processes, there has been little investigation into the long-term effects of nickel exposure. Our earlier studies on human lung epithelial cells showed that nickel exposure caused alterations to the expression of hundreds of genes, which did not revert to pre-exposure levels even after the termination of exposure [4]. Consequently, the cells underwent epithelial-mesenchymal transition (EMT), with the cells becoming invasive and migratory. The mesenchymal phenotype was persistent and continued even after the termination of Ni exposure [4]. EMT, the process during which the epithelial cells lose their polarity and cell-cell adhesion and acquire migratory and invasive phenotype, is associated with asthma, bronchitis, fibrosis, cancer and metastasis, diseases commonly associated with nickel exposure [1, 2, 5–7]. This suggests that the persistent transcriptional changes caused by nickel exposure could be a predisposing factor in the development of Ni-exposure-associated diseases. Therefore, unravelling the mechanisms that underlie the global gene expression changes that persist post nickel-exposure is critical for comprehensive understanding of the long-term impact of nickel exposure on human health.

In contrast to genotoxic metals, nickel does not form DNA adducts or DNA breaks and therefore is considered non-mutagenic [8–10]. Furthermore, oxidative stress caused by nickel exposure is significantly lower compared to other toxic metals [11]. Interestingly, a number of studies suggest that epigenetic alterations underlie nickel pathogenesis. Nickel ions inhibit histone acetyltransferases causing pervasive deacetylation [12–14]. Nickel also inhibits H3K9 demethylases resulting in increase in the levels of H3K9me2 [15]. In addition, nickel disrupts repressive chromatin domains marked by H3K9me2 and causes its spreading [16]. Increase in DNA methylation, has also been observed in nickel-exposed cells [17]. These findings suggest epigenome as an important target of nickel exposure. However, the genome wide dynamics of epigenetic alterations that occur post nickel-exposure, which potentially drives persistent changes to the transcriptome remains elusive.

To elucidate the mechanisms that underlie nickel induced long-term transcriptional aberrations, we examined the transcriptome and the epigenome of human lung epithelial cells during nickel exposure and after the termination of exposure. Transcriptional alterations are associated with histone modification signatures. Among the various histone modifications, H3K4me3, a histone modification associated with transcriptionally active genes is strongly predictive of expression changes [18]. Therefore, we generated global profiles of H3K4me3 to examine the epigenome during and after nickel exposure.

Our studies have identified a large number of genes whose expression was persistently altered by nickel exposure. Surprisingly, we found that the majority of gene expression changes occurred only after the termination of nickel exposure. Furthermore, we have identified substantial post-nickel-exposure alterations to the epigenome, which correlated with the observed gene expression changes. By identifying the epigenomic and transcriptomic changes that occur only after the termination of exposure, our studies shed new light on the persistent effects of nickel exposure and provide a new understanding of the potential long-term deleterious consequences of nickel exposure on human health.

## Results

### Nickel exposure induces persistent gene expression changes

BEAS-2B cells were exposed to a non-cytotoxic concentration of 100 µM NiCl_2_ for 6 weeks (nickel-exposed) [4]. After exposure, the cells were washed and plated in nickel-free medium at colony forming density and allowed to grow as colonies originating from individual nickel exposed cells. The colonies were then isolated and expanded as populations of nickel-washed-out cells for >6 months (Fig. 1a). Our earlier studies had shown that all nickel exposed cells displayed similar transcriptional profiles [4]. Therefore, we randomly selected a population of nickel-washed-out cells for a detailed examination. Gene expression analysis of the untreated cells (UT), nickel-exposed cells (Ni-E) and nickel-washed-out cells (Ni-W) cells were performed using RNA-Seq.

**Figure 1.**
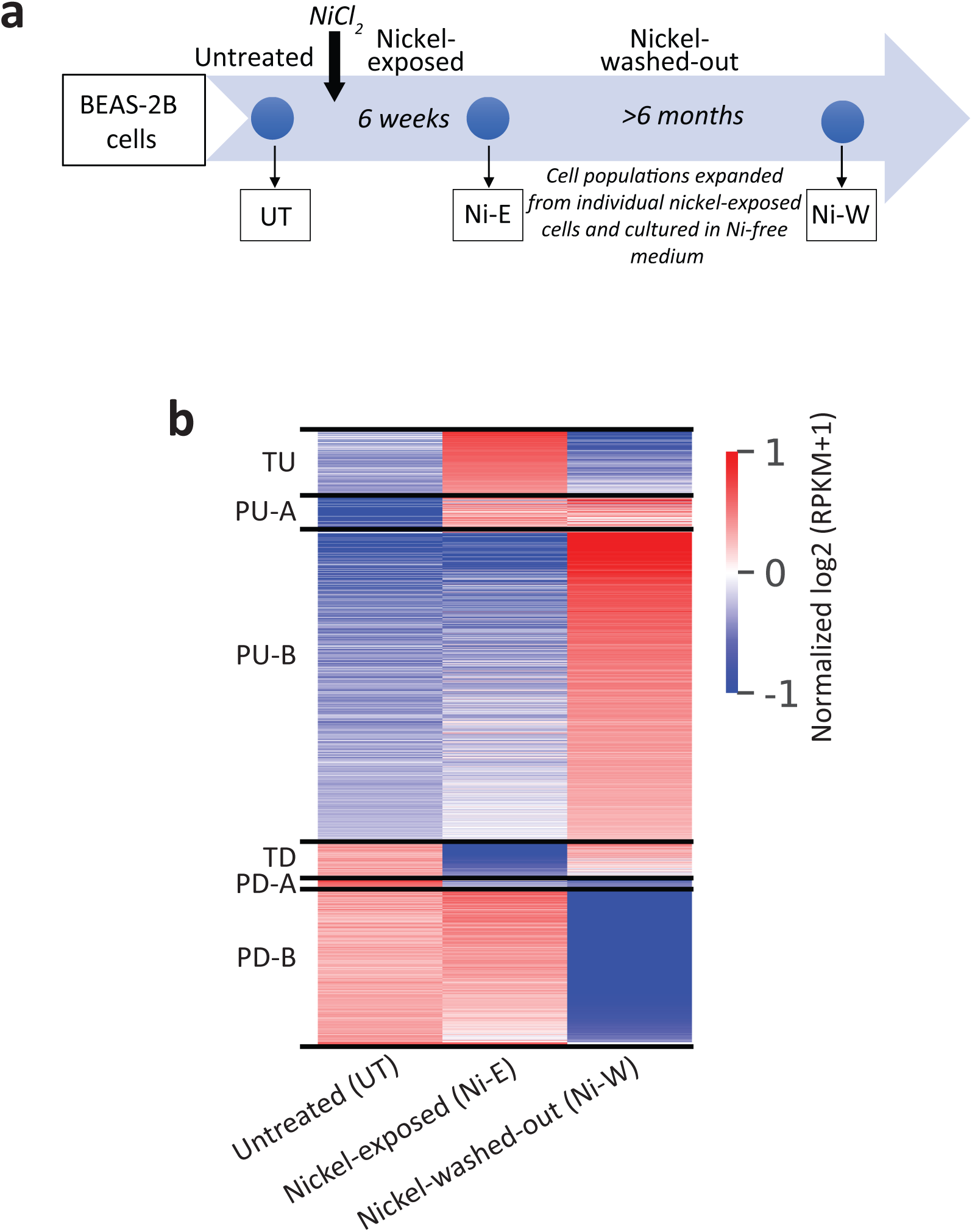
Nickel exposure causes persistent gene expression changes. **a** Schematic representation of nickel exposure. BEAS-2B cells were treated with 100 µM NiCl_2_ for 6 weeks (nickel-exposed). After exposure, the cells were plated at colony forming density to obtain populations of cells originating from individual nickel-exposed cells and grown in nickel-free medium for >6 months (nickel-washed-out). Gene expression was examined using RNA-Seq. **b** RNA-Seq results displayed as a heatmap showing the persistently and transiently differentially expressed genes (1.5 fold up-or down-regulation; Padj<0.05) in untreated (UT), nickel-exposed (Ni-E) and Ni-washed-out (Ni-W) cells. TU, transiently upregulated genes; PU-A, genes persistently upregulated in the presence of nickel; PU-B, genes persistently upregulated after the termination of exposure; TD, transiently downregulated genes; PD-A, genes persistently downregulated in the presence of nickel; PD-B, genes persistently downregulated after the termination of exposure. Normalized log2 transformed RPKM (pseudo count=1) values were used to generate the heatmap.

Examination of the differentially expressed genes revealed six gene-groups based on their expression profiles (Fig. 1b): ***i) Transiently upregulated (TU) genes:*** Nickel exposure upregulated 211 genes (in Ni-E cells), whose expression reverted to basal levels after the termination of exposure (in Ni-W cells); ***ii) Transiently downregulated (TD) genes:*** Nickel exposure downregulated 114 genes (in Ni-E cells), whose expression reverted to normal levels after the termination of exposure (in Ni-W cells). In addition to the transiently differentially expressed genes, we found a number of genes (1597 genes) that remained differentially expressed even after the cells were in culture for >6 months after the termination of nickel exposure. We termed these genes as persistently upregulated (PU) or persistently downregulated (PD). Interestingly, we identified two categories of PU genes: ***iii) Persistently upregulated-A (PU-A) genes:*** A subset of the PU genes (115 genes), which were upregulated during nickel exposure (in Ni-E cells) and the increased expression continued after the termination of exposure (in Ni-W cells). ***iv) Persistently upregulated-B (PU-B) genes***: A subset of PU genes (963 genes), whose expression remained unaltered during nickel exposure (in Ni-E cells). However, the genes were upregulated after the termination of exposure (Ni-W cells) and remained upregulated persistently. Similarly, the persistently downregulated genes could be classified into two categories: ***v) Persistently downregulated-A (PD-A) genes***: This set of genes were downregulated during nickel exposure (26 genes, in Ni-E cells) and continued to remain downregulated after the termination of exposure (in Ni-W cells); and ***vi) Persistently downregulated-B (PD-B) genes***: Expression levels of these genes were not altered during nickel exposure (in Ni-E cells). However, the genes were downregulated only after the termination of exposure (493 genes, in Ni-W cells).

### Nickel induces post-exposure genome-wide H3K4me3 changes

Next, we aimed at understanding the mechanisms underlying nickel-induced persistent gene expression alterations. Nickel is a non-mutagen and previous studies have shown that nickel exposure causes extensive changes to the epigenome [4, 16, 19]. Heritability of epigenetic changes through mitosis is well characterized. Therefore, it is conceivable that changes to the chromatin features could be an underlying cause for the nickel-induced gene expression changes that persist through cell division after the termination of exposure. To examine the relationship between nickel-induced gene expression changes and alterations to chromatin features, we profiled the genome-wide distribution of H3K4me3, a histone modification associated with transcriptional activation, in UT, Ni-E and Ni-W cells, using ChIP-Seq. We measured the levels of H3K4me3 at the gene promoters (+2 kb to −2 kb from transcription start site [TSS]) and quantified the changes in the H3K4me3 levels among the different categories of genes that we have identified (Fig. 1b) (see Methods for details).

#### Transiently upregulated (TU) genes

Nickel exposure transiently upregulated 211 genes. Examination of H3K4me3 at TU gene promoters showed no significant difference between Ni-E and UT cells (Fig. 2a-e). However, Ni-W cells showed significantly reduced H3K4me3 levels as compared to Ni-E and UT cells (Fig. 2a-e). This suggests that the TU gene upregulation in the presence of nickel is not accompanied by increase in H3K4me3 levels. However, the decrease in the expression of these genes after the termination of exposure is associated with loss of H3K4me3.

**Figure 2.**
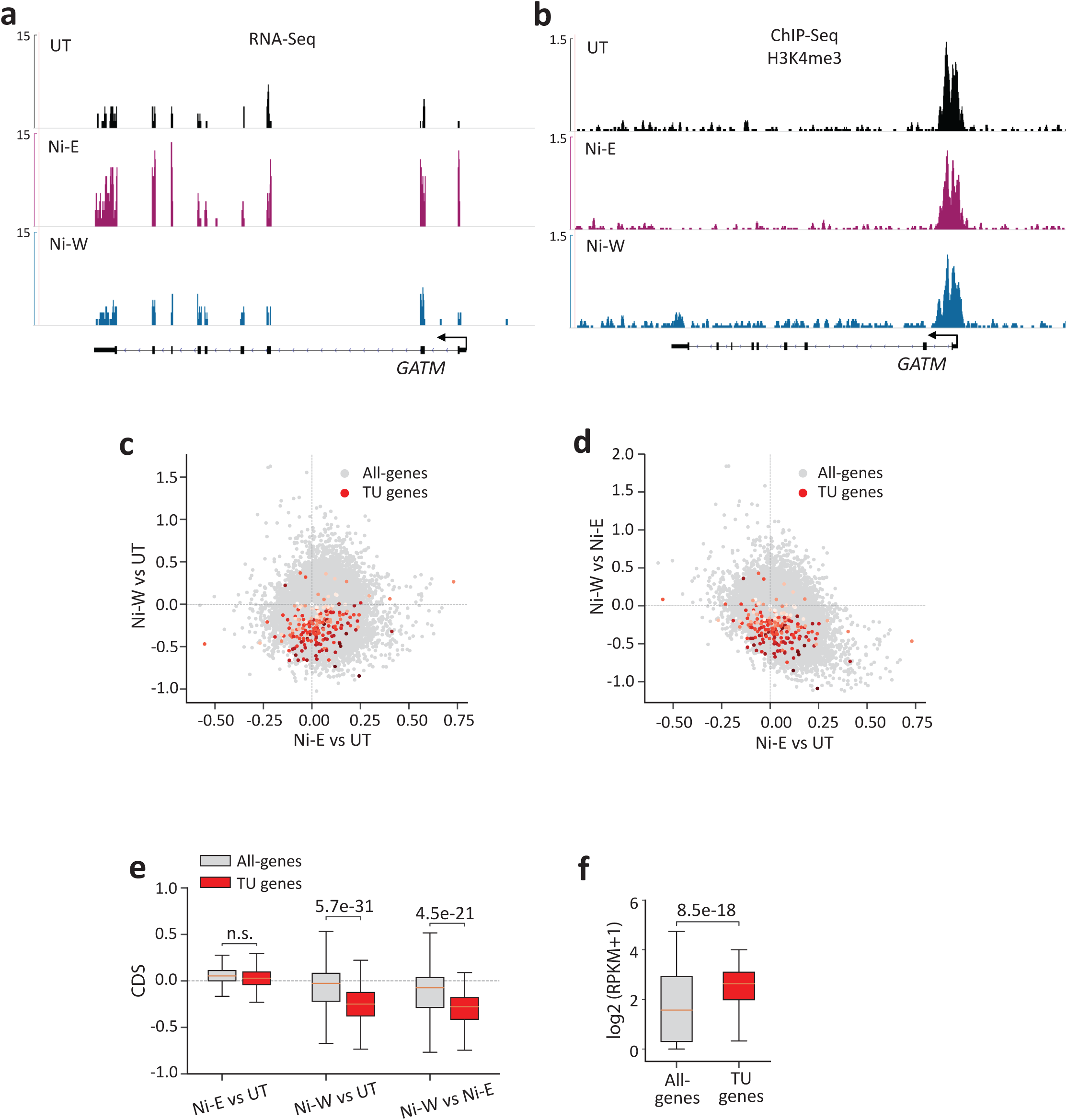
H3K4me3 profiles at the transiently upregulated (TU) genes. **a** Genome browser image showing expression levels (RNA-Seq) of *GATM*, a representative TU gene. **b** Genome browser image showing H3K4me3 at *GATM*. **c** and **d** Scatter plots showing changes in H3K4me3 levels at all the genes in the genome (all-genes) (light grey) and TU genes (red). x-axes in **c** and **d** show changes in the levels of H3K4me3 in Ni-E cells compared to UT cells. y-axis in **c** shows changes in the levels of H3K4me3 in Ni-W cells compared to UT cells, and y-axis in **d** shows changes in the levels of H3K4me3 in Ni-W cells compared to Ni-E cells. Intensity of red dots represent gene expression values in UT cells, with darker colors indicating higher expression. **e** Quantitation of the scatter plots **c** and **d**. **f** Box plot showing H3K4me3 levels at all-genes and TU genes. TU genes show higher H3K4me3 enrichment in untreated cells. P-values were calculated using Student’s t-test. n.s., not significant. TU (transiently upregulated), genes upregulated during nickel exposure, with the expression reverting to normal levels after the termination of exposure; CDS, chromatin dynamic score (see Methods for details).

Since the increase in the expression of TU genes was not associated with increase in H3K4me3 levels, we reasoned that that the genes that are upregulated in the presence of nickel could have permissive chromatin structure prior to nickel exposure. To examine this, we quantified the absolute H3K4me3 levels of all the gene promoters in the genome (all-genes) (+2 kb to −2 kb from TSS). We then compared the promoter H3K4me3 levels of TU genes and all-genes. Our analysis showed higher H3K4me3 levels at the promoters of TU genes compared to all-genes (Fig. 2f). Since H3K4me3 marks ‘open’ or accessible chromatin regions, our results suggest that the TU gene promoters exist in an accessible chromatin environment prior to nickel exposure.

#### Persistently upregulated (PU) genes

Nickel exposure persistently upregulated 1078 genes. Interestingly, while only 115 genes (10.7% of PU genes) were upregulated during nickel exposure (PU-A), 963 genes (89.3% of PU genes) were upregulated after the termination of exposure (PU-B). Examination of H3K4me3 at PU-A gene promoters did not reveal significant changes in H3K4me3 levels in Ni-E or Ni-W cells compared to UT cells (Fig. 3a-e). In addition, we did not find significant variations in H3K4me3 levels between Ni-E and Ni-W cells (Fig. 3a, b, d, e). This suggests that PU-A gene upregulation is not associated with increase in H3K4me3 levels. Interestingly, quantification of the H3K4me3 levels in untreated cells revealed higher enrichment H3K4me3 at the promoters of PU-A genes compared to all-genes (Fig. 3f). This suggests that PU-A gene promoters exist in a ‘open’ chromatin configuration prior to nickel exposure. PU-B gene promoters, on the other hand, showed significant increase in H3K4me3 levels in Ni-W cells compared to both UT and Ni-E cells (Fig. 3g-k). As expected, we did not detect H3K4me3 increase at PU-B gene promoters in Ni-E cells compared to UT cells (Fig. 3g, h, i, k), since this set of genes were upregulated only after the termination of nickel exposure (in Ni-W cells).

**Figure 3.**
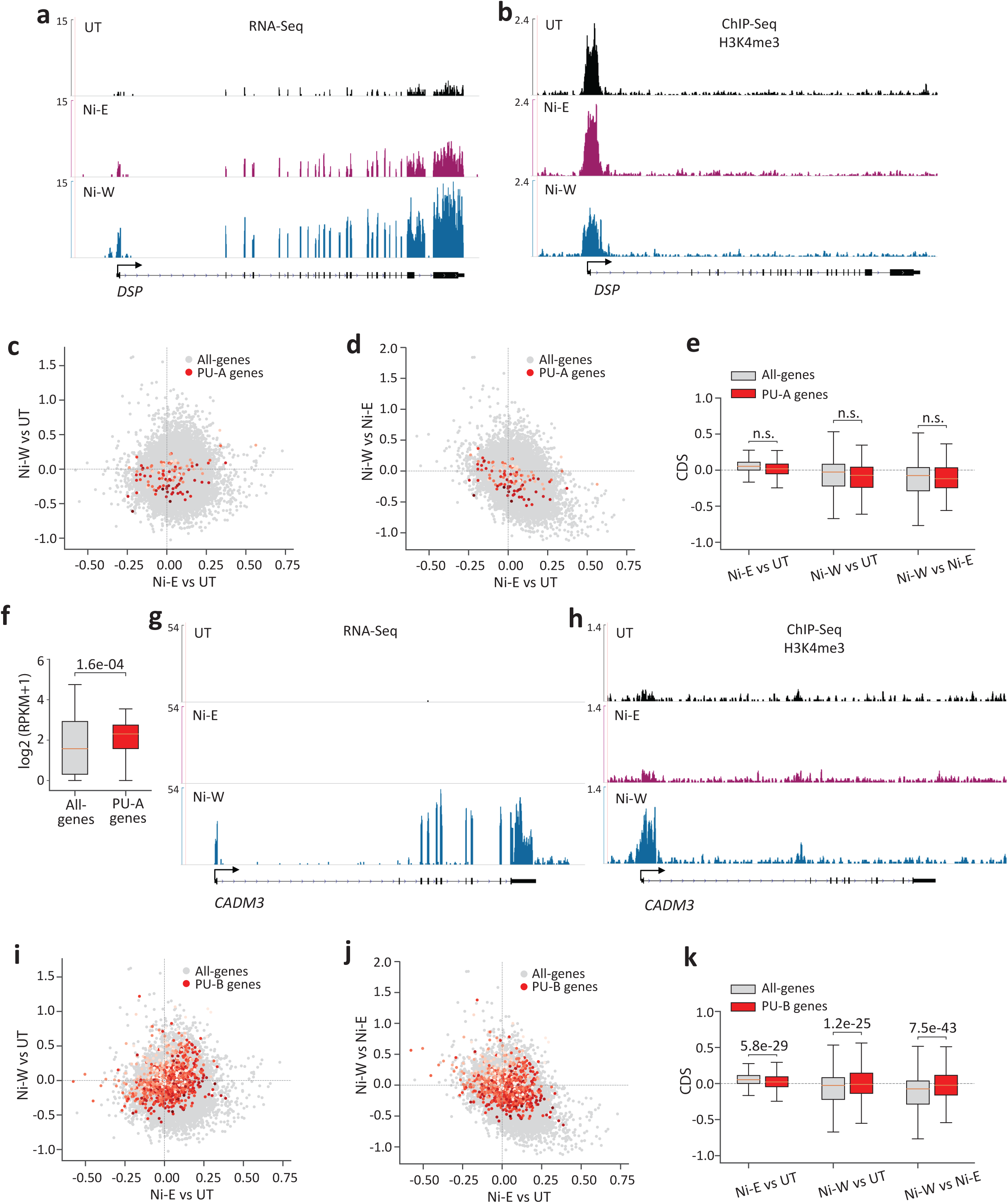
H3K4me3 profiles at persistently upregulated (PU) genes. **a** Genome browser image showing expression levels (RNA-Seq) of *DSP*, a representative PU-A gene. **b** Genome browser image showing H3K4me3 at *DSP*. **c** and **d** Scatter plots showing changes in H3K4me3 levels in all-genes (light grey) and PU-A genes (red). x-axes in **c** and **d** show changes in the levels of H3K4me3 in Ni-E cells compared to UT cells. y-axis in **c** shows changes in the levels of H3K4me3 in Ni-W cells compared to UT cells, and y-axis in **d** shows changes in the levels of H3K4me3 in Ni-W cells compared to Ni-E cells. Intensity of red dots represents the gene expression values in UT cells, with darker colors indicating higher expression values. **e** Quantitation of the scatter plots **c** and **d**. **f** Box plot showing H3K4me3 levels at all-genes and PU-A genes. PU-A genes show higher H3K4me3 enrichment in untreated cells. **g** Genome browser image showing expression levels (RNA-Seq) of *CADM3*, a representative PU-B gene. **h** Genome browser image showing H3K4me3 at *CADM3*. **i** and **j** Scatter plots showing changes in H3K4me3 levels in all-genes (light grey) and PU-B genes (red). x-axes in **i** and **j** show changes in the levels of H3K4me3 in Ni-E cells compared to UT cells. y-axis in **i** shows changes in the levels of H3K4me3 in Ni-W cells compared to UT cells, and y-axis in **j** shows changes in the levels of H3K4me3 in Ni-W cells compared to Ni-E cells. Intensity of red dots represent the gene expression values in UT cells, with darker colors indicating higher expression. **k** Quantitation of the scatter plots **i** and **j**. P-values were calculated using Student’s t-test. n.s., not significant. PU-A (persistently upregulated-A), genes upregulated during nickel exposure with the increased expression continuing after the termination of exposure; PU-B (persistently upregulated-B), genes upregulated after the termination of nickel exposure; CDS, chromatin dynamic score (see Methods for details).

Collectively, our results show that upregulation of gene expression in the presence of nickel (TU-A and PU-A) does not involve changes in H3K4me3 profiles. These genes possessed high levels of H3K4me3 levels prior to exposure (in UT cells), suggesting that they exist in an open chromatin environment in untreated cells. In contrast, the genes that were upregulated only after the termination of exposure (in Ni-W cells) were associated with significant increase in the levels of H3K4me3.

#### Transiently downregulated (TD) genes

Nickel exposure transiently downregulated 114 genes. Although we detected modest H3K4me3 loss in Ni-E and Ni-W cells compared to UT cells (Fig. 4a-d), these changes were not statistically significant (Fig. 4e), suggesting that both down regulation of gene expression in Ni-E cells and subsequent reversal of the downregulation in Ni-W cells were not associated with H3K4me3 changes.

**Figure 4.**
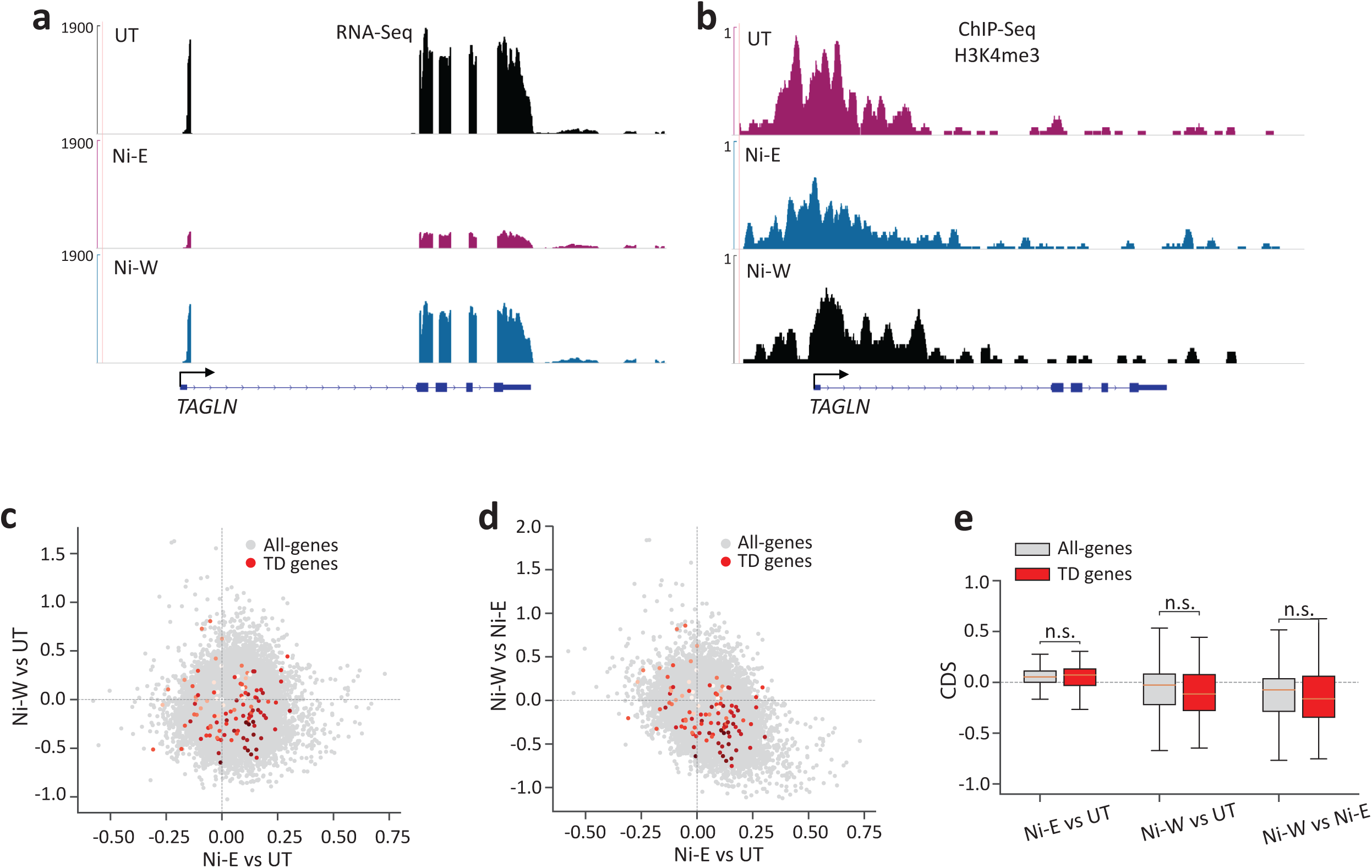
H3K4me3 profiles at the transiently downregulated (TD) genes. **a** Genome browser image showing expression levels (RNA-Seq) of *TAGLN*, a representative TD gene. **b** Genome browser image showing H3K4me3 at *TAGLN*. **c** and **d** Scatter plots showing changes in H3K4me3 levels in all-genes (light grey) and TD genes (red). x-axes in **c** and **d** show changes in the levels of H3K4me3 in Ni-E cells compared to UT cells. y-axis in **c** shows changes in the levels of H3K4me3 in Ni-W cells compared to UT cells, and y-axis in **d** shows changes in the levels of H3K4me3 in Ni-W cells compared to Ni-E cells. Intensity of red dots represents the gene expression values in UT cells, with darker colors indicating higher expression. **e** Quantitation of **c** and **d** *C* and *D*. P-values were calculated using Student’s t-test. n.s., not significant. TU (transiently downregulated), genes downregulated during nickel exposure, with the expression reverting to normal levels after the termination of exposure; CDS, chromatin dynamic score (see Methods for details).

#### Persistently downregulated (PD) genes

Nickel exposure persistently downregulated (PD) 519 genes. Interestingly, while merely 26 genes (5%) were downregulated during nickel exposure (PD-A), 493 genes (95%) were downregulated only after the termination of exposure (PD-B). Examination of PD-A gene promoters did not reveal changes in H3K4me3 levels in Ni-E and Ni-W cells as compared to UT cells (Fig. 5a-e). In contrast, we detected a robust loss of H3K4me3 at the PD-B gene promoters in Ni-W cells compared to both UT and Ni-E cells (Fig. 5f-j). These results suggest that while downregulation of gene expression that happen in the presence of nickel does not involve changes in H3K4me3, downregulation that occurs after the termination of exposure is associated with significant H3K4me3 loss. As anticipated, we did not detect decrease in H3K4me3 levels at PD-B gene promoters in Ni-E cells compared to UT cells (Fig. 5f, g, h, j), since the genes were downregulated only in Ni-W cells.

**Figure 5.**
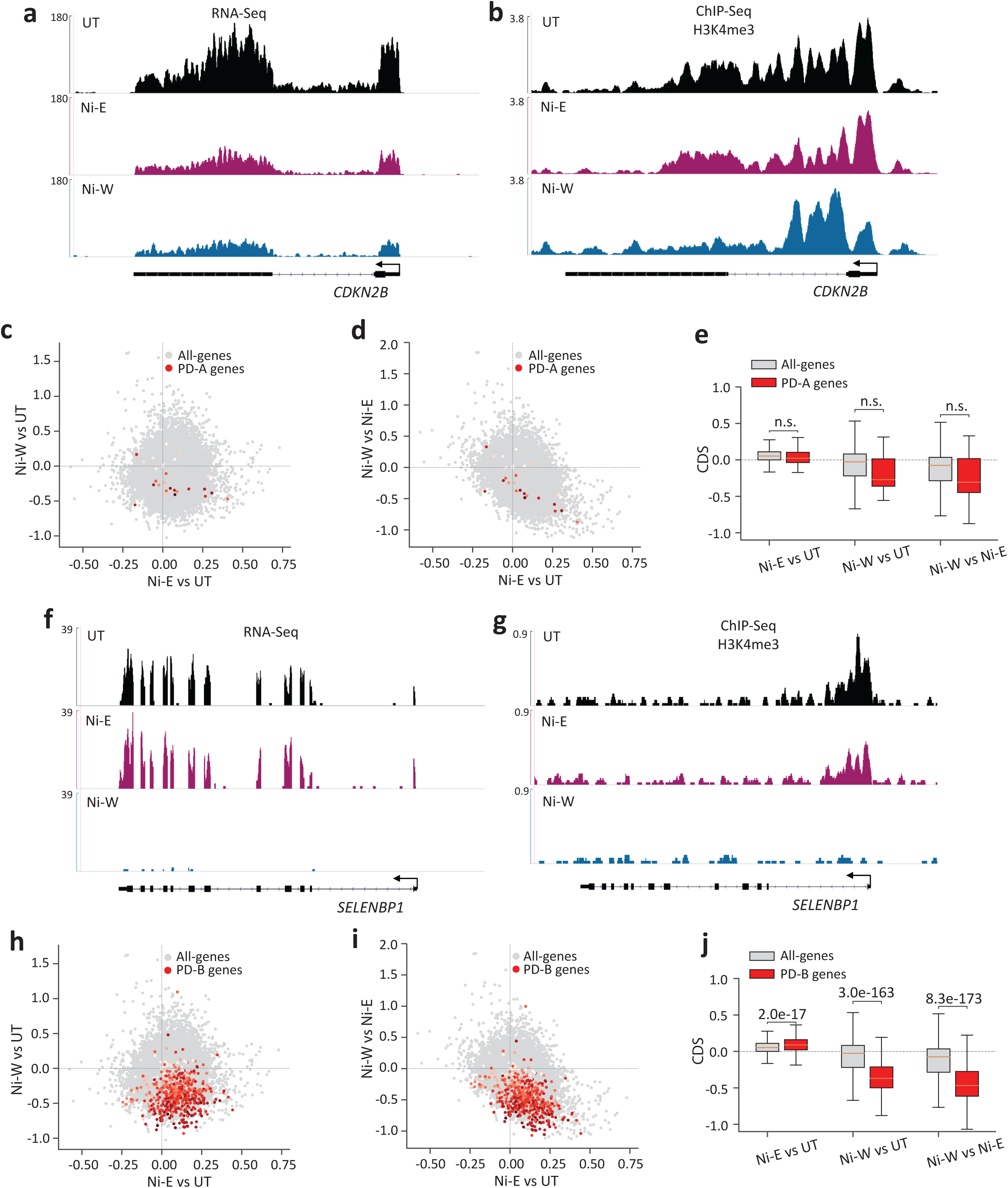
H3K4me3 profiles at the persistently downregulated (PD) genes. **a** Genome browser image showing expression levels (RNA-Seq) of *CDKN2B*, a representative PD-A gene. **b** Genome browser image showing H3K4me3 at *CDKN2B*. **c** and **d** Scatter plots showing changes in H3K4me3 levels in all-genes (light grey) and PD-A genes (red). x-axes in **c** and **d** show changes in the levels of H3K4me3 in Ni-E cells compared to UT cells. y-axis in **c** shows changes in the levels of H3K4me3 in Ni-W cells compared to UT cells, and y-axis in **d** shows changes in the levels of H3K4me3 in Ni-W cells compared to Ni-E cells. Intensity of red dots represents the gene expression values in UT cells, darker colors indicating higher expression values. **e** Quantitation of **c** and **d**. **f** Genome browser image showing expression levels (RNA-Seq) of *SELENBP1,* a representative PD-B gene. **g** Genome browser image showing H3K4me3 at *SELENBP1*. **h** and **i** Scatter plots showing changes in H3K4me3 levels in all-genes (light grey) and PD-B genes (red). x-axes in **h** and **i** show changes in the levels of H3K4me3 in Ni-E cells compared to UT cells. y-axis in **h** shows changes in the levels of H3K4me3 in Ni-W cells compared to UT cells, and y-axis in **i** shows changes in the levels of H3K4me3 in Ni-W cells compared to Ni-E cells. Intensity of red dots represent the gene expression values in UT cells, with darker colors indicating higher expression. **j** Quantitation of **h** and **i**. P-values were calculated using the Student’s t-test. n.s., not significant. PD-A (persistently downregulated-A), genes downregulated during nickel exposure with the downregulation continuing after the termination of exposure; PD-B (persistently downregulated-B), genes downregulated after the termination of nickel exposure; CDS, chromatin dynamic score (see Methods for details).

### Nickel-induced persistent gene upregulation is associated with loss of K3K27me3

Our results show that gene expression changes that occur in the presence of nickel were not associated with H3K4me3 alterations. Therefore, we next asked whether repressive histone modifications contribute to persistent transcriptional changes during nickel exposure. To accomplish this, using ChIP-qPCR, we examined the levels of repressive histone modification, H3K27me3, at the promoters of PU-A, PU-B, PD-A and PD-B genes. Our analyses detected significant decrease in the levels of H3K27me3 at PU-A (Fig. 6a) and PU-B (Fig. 6b) gene promoters in both Ni-W cells compared to UT cells. This suggested that H3K27me3 loss could underlie gene upregulation both in the presence of nickel and after the termination of exposure. Next, to understand whether the persistently downregulated genes gained repressive epigenetic marks, we examined H3K27me3 levels at the promoters of several PD-A and PD-B gene promoters. Our analysis did not reveal significant increase in the levels of H3K27me3 at the PD-A or PD-B gene promoters (Fig. 6c, d). These results suggest that persistent gene downregulation is not associated with the gain of repressive H3K27me3.

**Figure 6.**
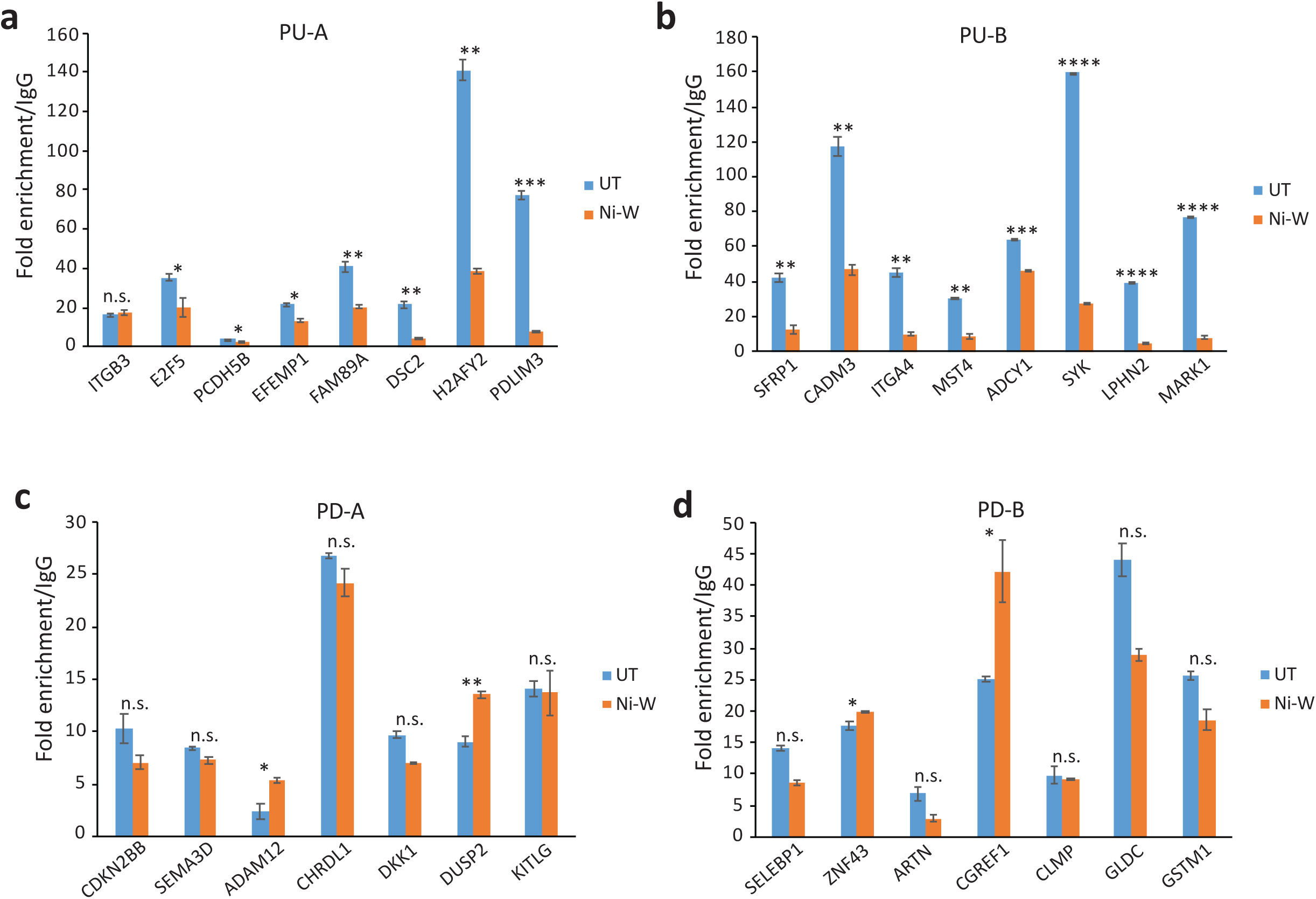
H3K27me3 is lost at the promoters of persistently upregulated (PU) genes. **a-d** ChIP-qPCR comparison of H3K27me3 levels between UT and Ni-W cells at the promoters of candidate PU-A (**a**), PU-B (**b**), PD-A (**c**) and PD-B (**d**) genes. While PU gene promoters showed loss of H3K27me3 (*A* and *B*), no significant alterations in H3K27me3 levels were detected at the PD gene promoters **c** and **d**. ChIP-qPCR results shown as fold enrichment of H3K27me3 over IgG. Error bars correspond to standard deviations from at least two biological replicates. In **a** and **b**, ‘*’ indicates statistically significant decrease in H3K27me3 levels in Ni-W cells compared to UT cells. In **c** and **d**, ‘*’ indicates statistically significant increase in H3K27me3 levels in Ni-W cells compared to UT cells. Statistical significance was evaluated using *t*-test (*P* < 0.05*; *P* < 0.01**; *P* < 0.001***; *P* < 0.0001****). n.s., not significant.

In summary, our results show that the up- and down-regulation of gene expression that occur during nickel exposure do not involve changes in the levels of the activating histone modification, H3K4me3. However, the gene expression changes that occur after the termination of exposure is associated with significant changes in H3K4me3. In contrast, the gene upregulation that occurred both in the presence of nickel and after the termination of exposure was associated with loss of repressive H3K27me3.

### Post-exposure transcriptional changes occur immediately after the termination of nickel exposure

In this study, the Ni-W cells that were used to identify post-exposure persistent transcriptional changes were in culture for >6 months after the termination of 6-week-Ni exposure (Fig. 1a). Hence, the post-nickel-exposure transcriptional changes that we have identified could have either occurred immediately after termination of nickel exposure or it could be the cumulative outcome of transcriptional changes in the cells over the course of >6 months. Therefore, to understand when the post-exposure transcriptional changes happened, we next compared the gene expression profiles of Ni-W cells (6-week nickel exposure followed by >6 months in culture without Ni) with that of Ni-W-2W cells (6-week nickel exposure followed by 2 weeks in culture without Ni) As shown in Figure 7, Ni-W and Ni-W-2W cells exhibit similar gene expression profiles, which is clearly different from that of the gene expression profile of Ni-E cells. These results suggest that the post-exposure persistent transcriptional changes that we have identified in Ni-W cells is established soon after the termination of nickel exposure and remained stable for >6 months.

**Figure 7.**
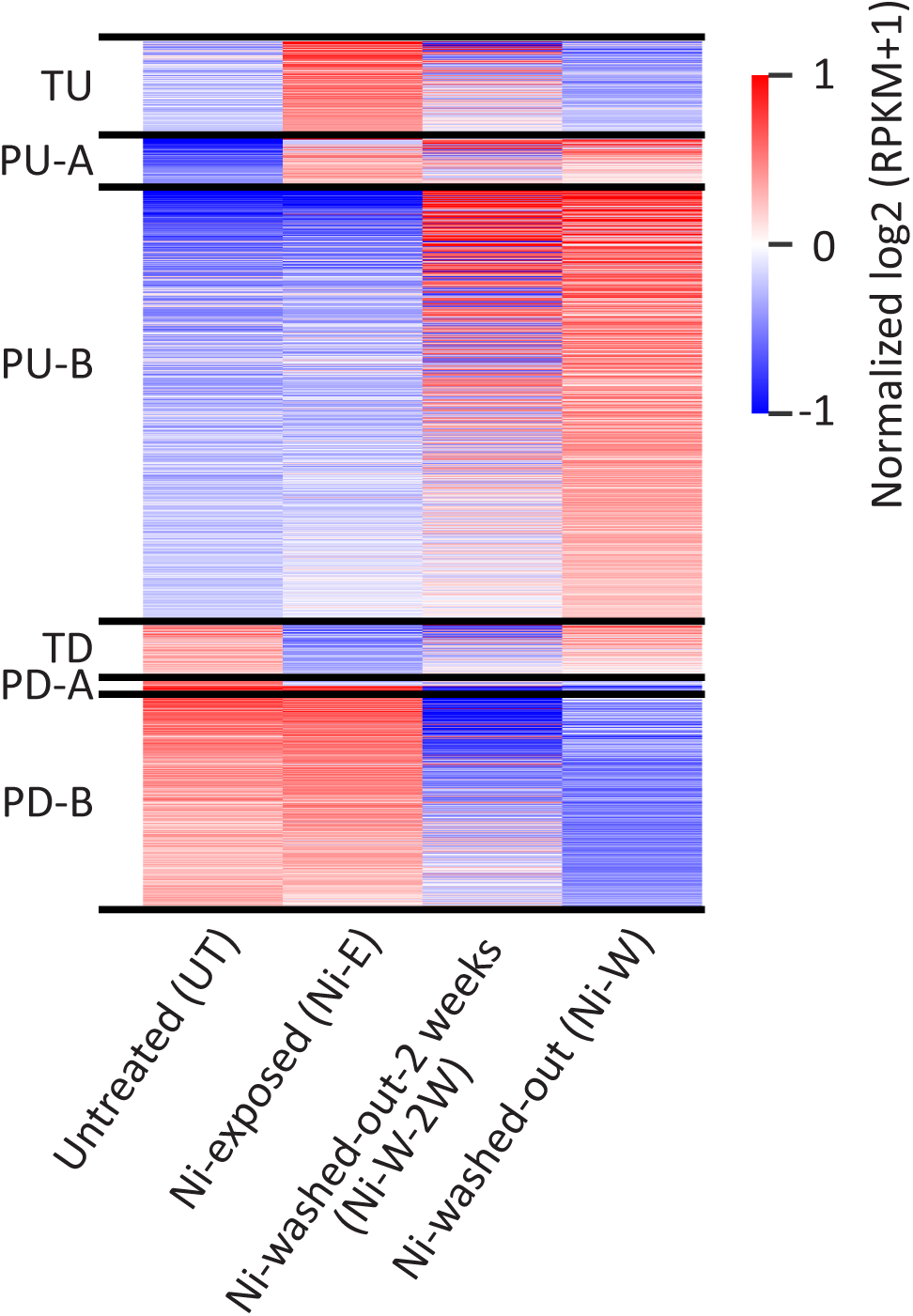
Post-nickel-exposure transcriptional changes in BEAS-2B cells occurred immediately after the termination of exposure. Following 6-week nickel exposure, the cells were grown in Ni-free medium i) for 2 weeks (Ni-W-2W) and ii) for >6 months (Ni-W). Gene expression was examined using RNA-Seq and the results are displayed as a heatmap showing the persistently and transiently differentially expressed genes (1.5-fold up- or down-regulation, Padj <0.05). Ni-W-2W and Ni-W cells display similar gene expression profiles suggesting that the post-nickel-exposure transcriptional changes occurred soon after the termination of exposure. TU, transiently upregulated; PU-A, persistently upregulated in the presence of nickel; PU-B, persistently upregulated after the termination of exposure; TD, transiently downregulated; PD-A, persistently downregulated in the presence of nickel; PD-B, persistently downregulated after the termination of exposure. Normalized log2 transformed RPKM (pseudo count=1) values were used to generate the heatmap.

### RT4 cells undergo transcriptional alterations after the termination of nickel exposure

Our results show that nickel could induce significant post-exposure effects in the lung epithelial BEAS-2B cells. We next asked whether non-lung cells undergo similar post-nickel-exposure transcriptional changes. To investigate this, we exposed human non-invasive bladder cancer RT4 cells to 100 µM NiCl_2_ for 6 weeks. After exposure, the cells were washed and plated in nickel-free medium for 2 weeks. Following this, gene expression analysis was carried out using RNA-Seq. As shown in Figure 8a, we identified the same six differentially expressed gene groups that we identified in BEAS-2B cells (Fig. 1b). This suggests that nickel exposure induces gene expression changes after the termination of exposure (PU-B and PD-B) in non-lung RT4 cells as well. However, relatively fewer genes were differentially expressed after the termination of nickel exposure in RT4 cells as compared to BEAS-2B cells.

**Figure 8.**
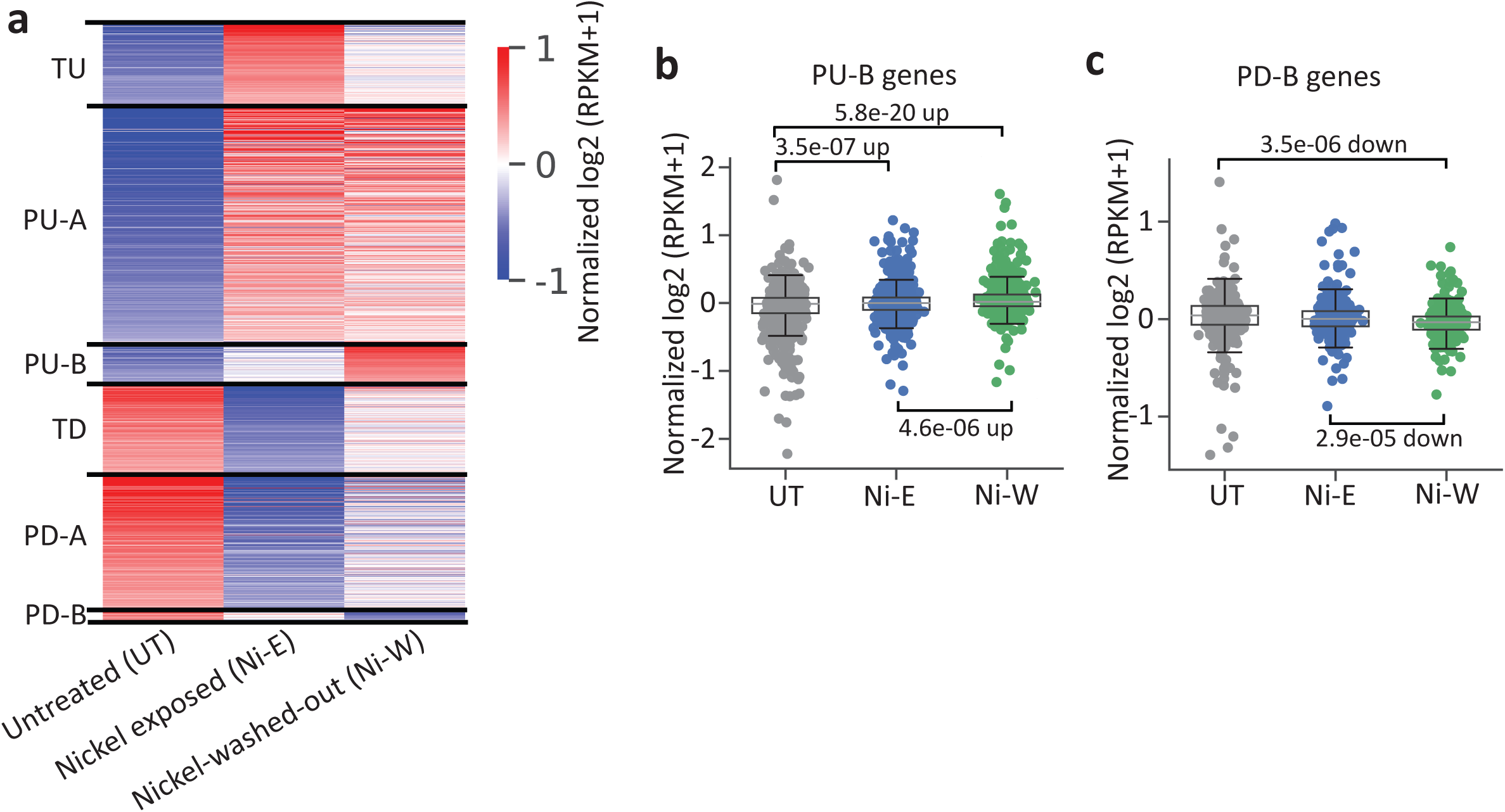
Nickel exposure induces persistent gene expression changes in RT4 cells. **a** Heatmap of RNA-Seq results in RT4 cells showing the persistently and transiently differentially expressed genes (1.5 fold up- or down-regulation; Padj<0.05) in untreated (UT), nickel exposed (Ni-E) and Ni-washed-out (Ni-W) cells. Normalized log2 transformed RPKM (pseudo count=1) values were used to generate the heatmap. TU, transiently upregulated; PU-A, persistently upregulated in the presence of nickel; PU-B, persistently upregulated after the termination of exposure; TD, transiently downregulated; PD-A, persistently downregulated in the presence of nickel; PD-B, persistently downregulated after the termination of exposure. **b** and **c** Normalized gene expression levels at the 3 conditions in RT4 cells. The PU-B (**b**) and PD-B (**c**) genes identified in BEAS-2B cells displayed similar patterns of differential expression in RT4 cells. Normalized log2 transformed RPKM (pseudo count=1) were used. P-values were calculated using Student’s t-test.

We next asked whether the genes that are differentially expressed after the termination of nickel exposure are conserved between BEAS-2B and RT4 cells. Surprisingly, our analyses showed that the PU-B and PD-B genes identified in BEAS-2B cells displayed similar patterns of differential expression in RT4 cells (Fig. 8b, c). This suggests that common upstream regulator(s) may regulate the genes that are differentially expressed after the termination of nickel exposure in different cell types.

## Discussion

In this study, by comprehensively examining the global gene expression profiles in lung epithelial cells during nickel exposure and after the termination of exposure, we report that majority of the nickel-induced transcriptional changes occur only after the termination of exposure. Furthermore, we show that the chromatin environment plays an important role in the temporality of gene expression changes caused by nickel exposure.

Genome-wide transcriptional changes caused by nickel exposure persist long after the termination of exposure [4]. Here, we identified two categories of persistently differentially expressed genes based on the timing of expression changes: i) the genes that were differentially expressed during nickel exposure (persistently upregulated-A [PU-A] and persistently downregulated-A [PD-A] genes); and ii) the genes that were differentially expressed only after the termination of nickel exposure (persistently upregulated-B [PU-B] and persistently downregulated-B [PD-B] genes) (Fig. 1). Interestingly, examination of the chromatin environment of the PU-A genes showed that these genes were enriched for both the activating histone modification, H3K4me3, and the repressive histone modification, H3K27me3, prior to nickel exposure. Enrichment of both activating and repressive histone modifications suggests bivalent chromatin at the PU-A gene promoters. Promoter bivalency typically maintains genes in a silent or poised-for-activation state [20]. Therefore, the PU-A genes exist in a poised chromatin environment prior to nickel exposure. Interestingly, Ni-induced upregulation of PU-A genes was associated with reduction in the levels of H3K27me3, although the H3K4me3 levels remained unchanged. This suggests bivalency resolution at PU-A gene promoters as a consequence of nickel exposure. Our earlier studies on the EMT master regulator gene, *ZEB1*, had indeed revealed nickel-induced bivalency resolution at its promoter, via loss of H3K27me3 [4]. The monovalent chromatin structure was irreversible even after the termination of exposure, resulting in the persistent activation of *ZEB1* [4]. Based on these results, it could be speculated that PU-A genes are predominantly bivalent, which are activated by the resolution of repressive bivalency through loss of H3K27me3, while the H3K4me3 levels remain constant. PU-B genes, on the other hand are largely devoid of H3K4me3 before nickel exposure, while being enriched for H3K27me3, suggesting that these genes exist in a repressive chromatin environment prior to nickel exposure. Significant increase in the levels of H3K4me3 occurs after the termination of exposure, which coincided with the increase in gene expression. In addition, loss of H3K27me3 in this group of genes likely contributes to gene upregulation.

Our data show that the genes that are upregulated in the presence of nickel (TU and PU-A genes) display high levels of H3K4me3 levels prior to nickel exposure (in UT cells). This suggests open or poised chromatin environment. On the other hand, upregulation of gene expression that occur only after the termination of nickel exposure (PU-B genes) is clearly associated with increase in H3K4me3 (in Ni-W cells). This suggests that these genes may exist in a relatively inaccessible chromatin conformation, and increase of H3K4me3 could be important for creating accessible chromatin and gene activation.

The persistently downregulated PD-A and PD-B genes, as expected, display high H3K4me3 prior to nickel exposure. Interestingly, while PD-A gene downregulation, which occurs in the presence of nickel was not associated with decrease in H3K4me3 levels, PD-B gene downregulation was significantly associated with H3K4me3 loss. Collectively our data suggest that the chromatin environment dictates the temporal dynamics of regulation of gene expression by nickel exposure.

Fascinatingly, our results show that the epigenome and transcriptome undergo robust changes after the termination of nickel exposure. The genes that undergo transcriptional changes in response to stimuli are classically categorized into immediate early or primary response genes and delayed or secondary response genes. The expression of secondary response genes, which may typically occur in a few hours or days after the initial stimulus, depend on translation of the primary response gene mRNAs and synthesis of proteins [21]. The PU-B and PD-B category genes however, could not be classified as delayed or secondary response genes since the cells were in culture in the presence of nickel for 6 weeks, and the expression changes happened only after the removal of nickel. Therefore, the mechanisms underlying the gene expression changes that occur after the termination of nickel exposure is likely different from that of the mechanisms that regulate secondary response genes.

Analysis of the sequences underlying the H3K4me3-gain regions (PU-B gene promoters) (Fig. 9a) and H3K4me3-loss regions (PD-B gene promoters) (Fig. 9b) to predict potential cis-regulatory elements suggest association of these loci with several chromatin regulators. These include known transcriptional activators such as MED, BRD2, BCL3 and E2F4 and transcriptional repressors such as REST, JARID2, MBD3 and MECP2, suggesting that these loci are hotspots of dynamic chromatin changes. However, the mechanisms through which nickel exposure impacts these regions to cause changes after the termination of exposure is still unclear. Interestingly, our results show that the genes that are differentially expressed after the termination of nickel exposure are shared between BEAS-2B (lung) and RT4 (bladder) cells (Fig. 8). Similar genes being affected in two very different cell types after the termination of exposure suggests that the post-exposure effects could be controlled by common regulatory mechanisms.

**Figure 9.**
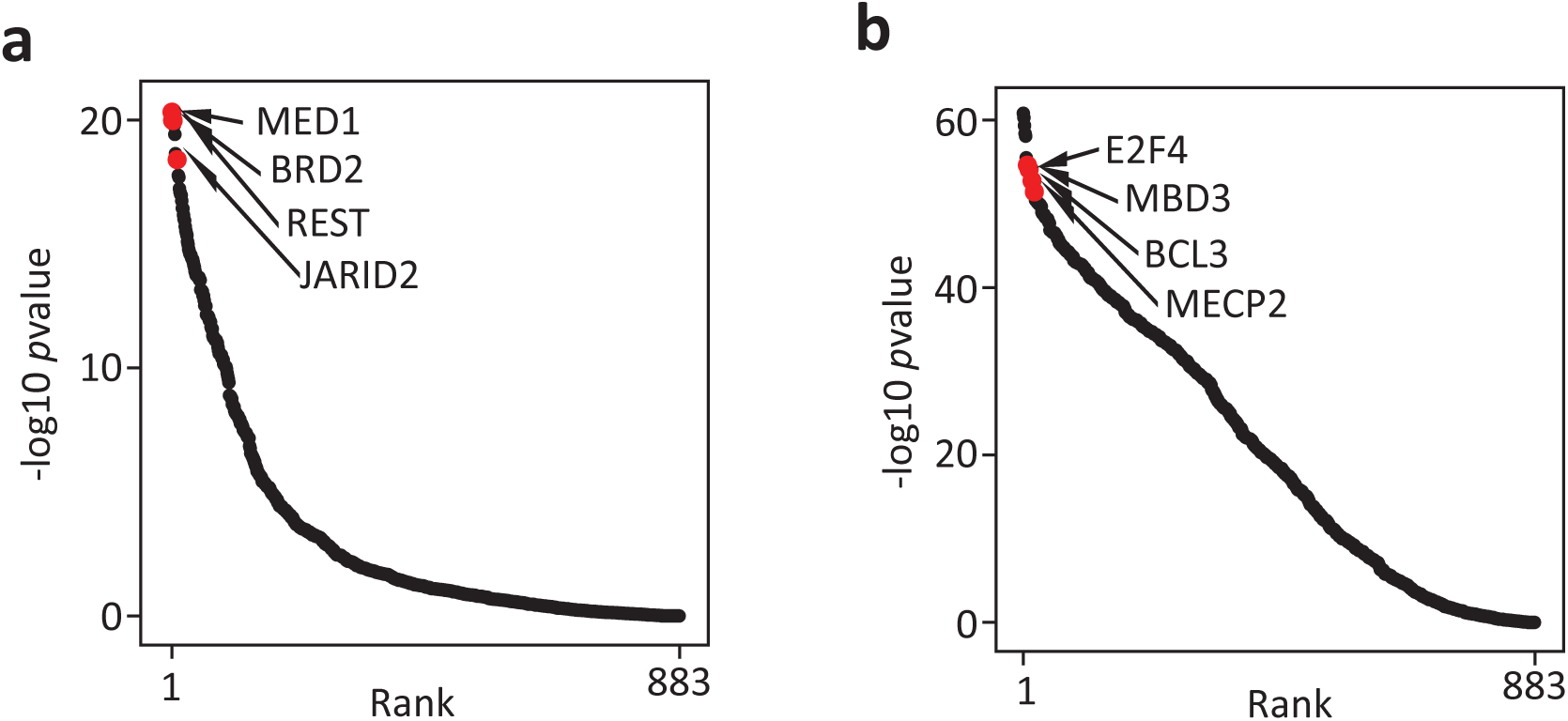
Differential-H3K4me3 regions at the PU-B and PD-B gene promoters potentially bind chromatin regulators. TF binding is predicted using BART and ranked by P-value scores. Top tanked factors are predicted as high-confident factors that bind the query regions. **a** TF enrichment at PU-B genes promoters, which display increased H3K4me3 levels. **b** TF enrichment at PD-B gene promoters, which display decreased H3K4me3 levels.

Nickel ions are known to affect protein activities in various ways. For example, nickel inhibits the HIF proline hydroxylases and stabilizes the HIF protein thereby inducing hypoxia-like state in the cells even under normoxia [22]. Several Jumonji C (JmjC) domain containing demethylases (JHDMs), which are iron and 2-oxoglutarate-dependent enzymes have hypoxia response elements (HREs) in their promoters and are activated by HIF-1 [23]. Therefore, nickel exposure could activate these enzymes. However, the JHDMs bind iron at their catalytic center [24] and nickel inhibits this family of enzymes by displacing the iron and impedes the histone demethylation process [25, 26]. Therefore, while nickel can increase the expression JmjC domain containing histone demethylases by stabilizing HIF, it can also inhibit their enzymatic activities by replacing iron at their catalytic centers. Similarly, nickel can substitute zinc at the DNA binding domains of several zinc finger proteins, resulting in inhibition of DNA binding as well as alteration of DNA-binding specificity [16, 27]. Therefore, it is tempting to speculate that nickel exposure could initiate a first wave of transcription, during which the genes transcriptionally upregulated in the presence of nickel (PU-A genes) could potentially code for transcription factors such as chromatin regulators and master regulators of transcription (represented by Gene X and Protein X in Figure 10). However, the presence of nickel ions could potentially maintain these proteins inactive by inhibiting their enzymatic activities and/or by altering their DNA binding affinities. Termination of nickel exposure could render these proteins active, which leads to epigenomic alterations and target gene activation, thus initiating a second wave of differential expression, post exposure (represented by Gene Y and Protein Y in Figure 10). Therefore, the post-exposure alterations to the epigenome and transcriptome could be an outcome of this second wave of transcription for which removal of nickel may be essential. However, a thorough investigation is required to understand the mechanisms underlying the post-exposure effects of nickel.

**Figure 10.**
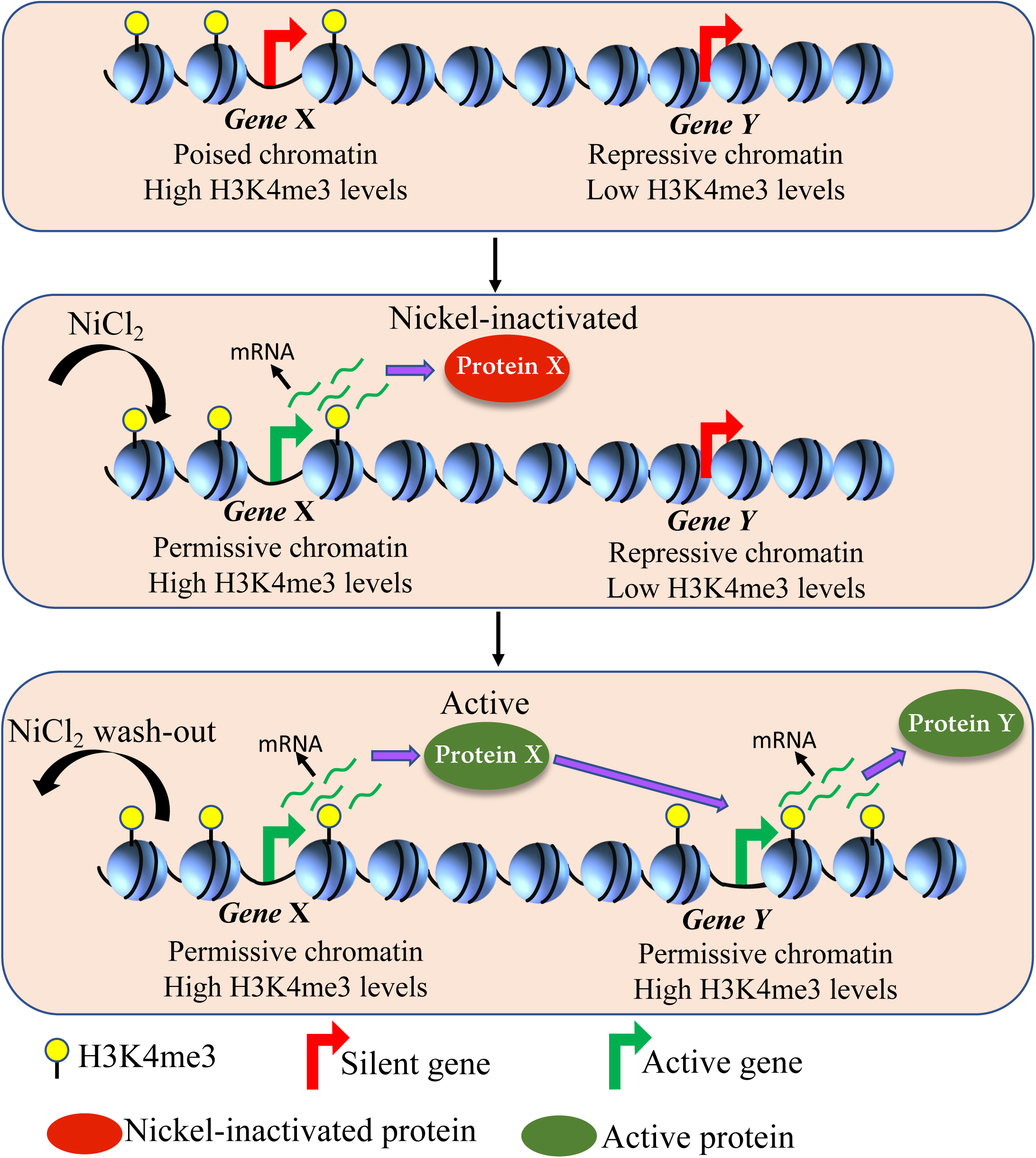
Proposed model for the post-nickel-exposure transcriptional changes. **a** In the untreated cells, Gene X, a potential chromatin regulator, is transcriptionally silent. However, Gene X promoter is enriched for H3K4me3 and possibly exists in a poised chromatin environment. Gene Y, also a transcriptionally silent gene, has low H3K4me3 levels and exists in a repressive chromatin environment. **b** Nickel exposure induces the first wave of transcription, where it persistently activates Gene X, probably through induction of an upstream transcriptional regulator and/or through resolution of bivalency by inducing loss of repressive chromatin modifications. However, the protein encoded by Gene X (Protein X [red]) remains inactive in the presence of Ni^2+^ ions either due to inhibition of its enzymatic activity or through alteration of its DNA-binding affinity. **c** Upon Ni wash-out, Protein X [green] is rendered active and initiates a second wave of transcription by altering the chromatin environment of Gene Y, which gains H3K4me3 and is transcriptionally activated (Protein Y).

Traditionally, toxicological studies are carried out during active exposure. However, the importance of persistent changes to the transcriptome is increasingly being realized. Early-life exposure to environmental pollutants such as heavy metals, pesticides and polycyclic aromatic compounds is implicated in disease susceptibility later in life [28, 29]. Moreover, smoking induced changes to expression of several genes, including oncogenes, tumor suppressor genes, and miRNAs failed to revert to never-smoker levels years after cessation of smoking, which could explain the persistent risk of lung cancer in former smokers [30–32]. These studies suggest that exposure-induced long-term changes could be a significant factor in the initiation and progression of environmental diseases. While these previous studies have shown irreversible effects of environmental exposures, our current study has uncovered a new category of transcriptional and epigenetic changes, which occur only after the termination of exposure. It is interesting that most of the transcriptomic and epigenomic changes occur only after the termination of nickel exposure. This highlights the importance of investigating the genome-wide temporal dynamics of environmental exposures to obtain a comprehensive understanding of adverse health outcomes.

## Methods

### Cell culture and treatments

Human lung epithelial BEAS-2B cells (ATCC, CRL-9609) were cultured in Dulbecco’s Modified Eagle’s Medium (DMEM, Cellgro) supplemented with 1% Penicillin-Streptomycin and 10% Fetal Bovine Serum (FBS, Atlanta Biologicals) at 37°C and 5 % CO_2_. For nickel exposure, 100 µM NiCl_2_ (N6136, Sigma) was added to the media and the cells were cultured for 6 weeks (Ni-exposed, Ni-E). The cells were split at 80% confluence. Cell culture medium was changed every 48 h and NiCl_2_ was added to the cells. Following 6-week exposure, a homogenous population of nickel-exposed cells was isolated as described earlier [4]. Briefly, the nickel-exposed cells were then plated in 15 cm plates at the rate of 1000, 500, 100 and 50 cells per plate in nickel-free medium. Nicely separated single colonies were randomly isolated and the populations were expanded in nickel-free medium. A randomly selected homogenous population of nickel-washed-out cells (Ni-W) in culture for >6 months post nickel-exposure (Ni-W cells) was used for a detailed examination. For short term nickel-wash-out, the nickel-exposed cells (100 µM NiCl_2_ for 6 weeks) were cultured in nickel-free medium for 2 weeks (Ni-W-2-week).

Human urinary bladder epithelial RT4 cells (a generous gift from Dr. Tang, New York University School of Medicine, NY) were cultured in McCoy’s 5A Modified Medium (Life Technologies, Carlsbad, CA), supplemented with 10% FBS (Atlanta Biologicals) and 0.5% Penicillin-Streptomycin at 37°C and 5% CO_2_. The cells were exposed to 100 µM NiCl_2_ for 6 weeks (Ni-E). Following exposure, the cells were washed and cultured in nickel-free medium for 2 weeks to obtain nickel-washed-out cells (Ni-W).

### RNA isolation, RNA-Seq and data analysis

Total RNA was isolated using RNeasy Kit (Qiagen 74104). RNA-Seq was performed using RNA samples isolated from two biological replicates. RNA-Seq libraries were prepared using Illumina TruSeq RNA Sample Preparation Kit (RS-122-2002) according to manufacturer’s protocol. Sequencing was performed using Illumina HiSeq 4000. RNA-Seq data analysis was performed using BioWardrobe Experiment Management System[33]. Briefly, the raw Fastq sequence files were aligned to the human genome (hg19) using STAR[34] (version 2.4.oj using default parameters; multi hits removed) with a known reference annotation gtf file from RefSeq. Gene expression levels were quantified as Reads Per Kilobase of transcript per Million mapped reads (RPKM) using BioWardrobe algorithm[33]. Differential gene expression was calculated using DESeq2[35]. Genes that showed ≥1.5 fold up- or down-regulation between the conditions compared, along with adjusted p-value<0.05 were considered as differentially expressed. To generate heatmaps, the log2 transformed RPKM values (pseudo count = 1) were first median-centered for each sample, and then mean-centered for each gene. Genes were decreasingly ranked by expression variance across different treatments (UT, Ni-E and Ni-W) within each group. The RNA-Seq data were deposited in the Gene Expression Omnibus (GEO) under the accession number GSE136558. RNA-Seq data of BEAS-2B nickel-washed-out-2 weeks (Ni-W-2W) cells was obtained from our earlier study (GSE95180) [4].

### Chromatin immunoprecipitation (ChIP) and ChIP-qPCR

ChIP was performed using samples isolated from two biological replicates. The cells were crosslinked with 1% formaldehyde for 10 minutes at 25°C and sonicated to obtain 200-500 bp fragments. ChIP was performed as described earlier [4, 16] using ChIP grade antibodies against H3K4me3 (Millipore, 07-473) and H3K27me3 (Millipore, 07-449). H3K27me3 ChIP DNA was analyzed using quantitative PCR (qPCR) analysis using FastStart Universal SYBR Green Master Mix (Roche) on a 7900HT Fast Real-Time PCR system (Applied Biosystems). Statistical significance of qPCR results was evaluated using t-test.

### ChIP-Seq and data analysis

H3K4me3 ChIP-Seq libraries were prepared using Illumina TruSeq ChIP sample preparation Kit (IP-202-1024), according to manufacturer’s protocol. Sequencing was performed Illumina HiSeq 4000 to obtain 50-nucleotide single-end reads. ChIP-Seq reads were aligned to the human reference genome (GRCH38/hg38) using BWA [36]. SAM files were then converted into BAM format using samtools [37]. Bedtools was used to convert BAM files into BED format. Only 1 copy of the redundant reads was retained. All non-redundant reads were included for downstream analyses without peak calling. ChIP-Seq data were deposited in the Gene Expression Omnibus (GEO) under the accession number GSE136558.

### Comparison of H3K4me3 levels between different gene categories

To examine the impact of nickel exposure on the different categories of genes that we identified, we quantified the H3K4me3 levels by counting the normalized ChIP-Seq reads (RPKM) at each gene’s promoter (+/-2kb from TSS). A chromatin dynamic score (CDS) was used to quantify the changes of histone modification levels for each gene *i* using the formula previously described[38, 39], and to compare treatment (t) vs control (c): 1. Ni-E vs UT; 2. Ni-W vs UT; and 3. Ni-W vs Ni-E.

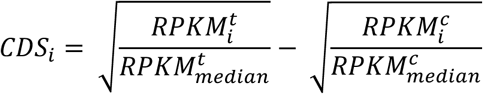

### Transcription factor binding prediction

A revised version of the BART algorithm [40] was used for transcription factor (TF) enrichment analysis. Briefly, a collection of union DNase I hypersensitive sites (UDHS) was previously curated as a repertoire of all candidate cis-regulatory elements in the human genome [41]. A total of 7032 ChIP-Seq datasets were collected for 883 TFs, with each TF having one or more ChIP-Seq datasets from different cell types or conditions. A binary profile was generated for each TF on UDHS, indicating whether the TF has a ChIP-Seq peak overlapping each UDHS. TF enrichment analysis was then applied to each TF by comparing the bindings on the UDHS overlapped promoter regions of selected genes and the UDHS overlapped promoter regions of all genes. P-value was obtained using the Fisher’s exact test for each TF. All TFs included in the analysis were ranked by P-value scores (-log10 P-value). Top ranked factors are predicted as high-confident factors that bind at the query regions.

## Author Contributions

S.C, C.C.J and C.Z designed the experiments. C.C.J, X.Z and SC performed the experiments. C.Z and S.C directed data analysis. Z.W, V.S.T and C.Z performed data analysis. S.C wrote the manuscript with contributions from C.C.J, Z.W and C.Z. All authors read and approved the final manuscript.

## Acknowledgements

This work was supported by National Institutes of Health (NIH) grants R01ES024727 and P30ES000260 pilot project to S.C., and K22CA204439 and R35GM133712 to C.Z. Research reported in this publication includes work performed in the NYUMC Genome Technology Center, partially supported by the Cancer Center Support Grant, P30CA016087, at the Laura and Isaac Perlmutter Cancer Center.

